# Damaged motile cilia undergo phagolysosomal clearance during regeneration

**DOI:** 10.64898/2026.03.31.715518

**Authors:** Muyang Ren, Kate Heesom, Charlotte Melia, Shinya Matsuda, Kentaro Nakano, Girish R. Mali

**Author notes:** Correspondence should be addressed to G.R.M.

## Abstract

The remodelling and regeneration of motile cilia are widespread yet poorly understood cellular responses to stress. Whether components from damaged cilia are reutilised during regeneration remains unclear. Here, we apply acute, non-lethal stress to the multiciliated eukaryote *Tetrahymena thermophila* to investigate the fate of damaged motile cilia during regeneration. We find that damaged axonemes accumulate intracellularly as coiled structures, termed ‘c-rings’, which are sequestered within phagosome-associated membrane compartments and subsequently associate with acidic compartments for clearance. Imaging, quantitative proteomics and genetic perturbations reveal a phagolysosomal route for processing damaged axonemes, accompanied by recruitment of autophagy-associated machinery. Remarkably, ciliary regeneration proceeds without detectable c-ring accumulation, even when new protein synthesis is inhibited, indicating that reutilisation of internalised axonemal material is not essential for ciliogenesis. Our findings uncover an unexpected mechanism for clearing damaged motile cilia and reveal how ciliary degradation and regeneration can operate as distinct cellular processes.

---

Cilia and flagella are surface-exposed extensions of the eukaryotic cytoskeleton that undergo extensive remodelling during development, cellular adaptation and stress. Mammalian cells shed or resorb their primary cilia during the cell cycle, epithelial morphogenesis and in response to environmental stress ^1–4^. Unicellular protists also employ diverse mechanisms of ciliary resorption and intracellular disassembly during their life cycles or following heat, hydrodynamic and osmotic stress ^5–7^. However, the turnover of multiple motile cilia, which form large and structurally complex cytoskeletal assemblies, remains poorly understood. Notably, mammalian respiratory epithelial cells exhibit intracellular accumulation of ciliary axonemes and basal bodies following coronavirus infection, alongside extracellular ciliary shedding ^8–10^. These observations raise a fundamental question: how do cells process and dispose of damaged motile cilia once their axonemal structures are displaced from the cell surface?

The fate of damaged ciliary material also has important implications for ciliary regeneration. Classical experiments in the biflagellate alga *Chlamydomonas reinhardtii* established the possibility that components released during ciliary resorption can be reutilised to assemble new cilia^11^. This model has influenced our understanding of ciliary regeneration, in which salvaged components are proposed to supplement a pre-existing cytoplasmic pool of ciliary precursors. However, whether such structural reutilisation occurs in multiciliated eukaryotes remains unclear. An alternative model proposes that ciliary resorption mobilises pre-existing precursor pools to initiate ciliogenesis independently of the direct reutilisation of old axonemal components ^12–14^. Crucially, the intracellular fate of damaged motile axonemes and their potential contribution to newly assembled cilia have not been directly resolved.

Here, using the multiciliated eukaryote *Tetrahymena thermophila*, we uncover an unexpected route for the intracellular clearance of damaged motile cilia following acute cellular stress. Combining fluorescence and volume electron microscopy, quantitative proteomics, live-cell tracking of ciliary proteins and genetic perturbations, we show that damaged axonemes accumulate as coiled intracellular structures that are sequestered within phagosomes and delivered to acidic compartments for clearance. These compartments are associated with autophagy-related machinery, suggesting a potential interface between phagocytic and autophagic pathways during the processing of damaged axonemes. Remarkably, ciliary regeneration proceeds independently of the accumulation of these structures, even when new protein synthesis is inhibited, supporting a model in which pre-existing precursor pools contribute to rebuilding motile cilia. Together, our findings reveal a previously unrecognised route for damaged axoneme clearance and suggest that ciliary degradation and regeneration can operate as distinct cellular processes to maintain organelle homeostasis.

## Results

### Stress-induced deciliation generates intracellular coiled axonemes

To study the intracellular processing of multiple motile cilia, we applied an acute, non-lethal calcium shock to the protozoan ciliate *Tetrahymena thermophila*. Dibucaine hydrochloride or controlled reduction of the cell’s pH in the presence of CaCl_2_ coupled with gentle mechanical shearing are well-established methods of deciliating *Tetrahymena*. Both induce complete shedding of surface cilia, the latter providing a non-lethal means to track the dynamics of ciliary regeneration in live cells ^15–17^ (**Fig.1a, b**). We found that omitting mechanical shearing elicits a partial de-ciliation response where over 75% of cilia are shed in line with previous work ^13^ (**Fig.1b**). High calcium paralyses the remaining locomotory and oral cilia by impacting motors and is known to destabilise microtubules ^18–22^. These paralysed cilia are observed on cell surfaces 30 seconds after deciliation and have previously been reported to be resorbed within the first 15-30 minutes after treatment in *Tetrahymena* (**Fig.1a, b)** ^13^. We observed acetylated ɑ-tubulin-positive coiled structures distributed throughout the cytoplasm of partially-deciliated cells fixed 30 minutes after the CaCl_2_/pH shock (**Fig.1c**). *Tetrahymena* cilia on the surface are 5-7 µm long (**Fig.1c, left panels**). The arc lengths of the coils broadly matched these dimensions, consistent with the intracellular accumulation of structurally preserved ciliary axonemes (**Fig.1c, right panels**). We refer to these coiled cilia as c-rings and propose that they originate from damaged surface cilia undergoing resorption. Quantification showed a significant increase in the frequency of c-rings relative to non-treated control cells (**Fig.1d**). Fully deciliated cells do not accumulate c-rings. Together, these observations indicate that partial deciliation is accompanied by the accumulation of highly coiled axonemal structures within the cytoplasm.

### Intracellular axoneme clearance accompanies ciliary regeneration

Fully or partially deciliated *Tetrahymena* undergo ciliary regeneration and regain partial motility by 90 minutes ^15^. Time-course immunostaining and quantification showed that recovery of deciliated cells to a fully ciliated state with restoration of ciliary lengths up to 7 µm and concomitant clearance of c-rings takes up to 240 minutes (**Fig.1e-g)**. C-rings are detected throughout the cytoplasm at 30 minutes post-treatment in cells that have no external cilia. Gradual recovery of cortical and oral apparatus cilia was observed at 120 minutes post-treatment with few c-rings persisting in the cytoplasm. C-rings are cleared from cells that had fully regenerated cilia by 240 minutes post-treatment. C-ring clearance and the re-emergence of new cilia occur simultaneously indicating that reutilisation of components is potentially possible but remains a correlative finding (**Fig.1g**). Alternatively, ciliary regeneration and c-ring clearance could be purely co-incidental yet independent cellular events.

### C-rings retain characteristic axonemal tubulin poly-modifications

The persistence of intracellular c-rings following partial deciliation raised the question of whether these structures retain molecular features characteristic of mature ciliary axonemes. Ciliary axonemes are enriched in tubulin post-translational modifications (PTMs) that influence microtubule stability and interactions with microtubule-associated proteins, including severing enzymes and depolymerising motors. We therefore used ultrastructure expansion microscopy (U-ExM) to examine whether c-rings retain the characteristic tubulin modifications of mature ciliary axonemes. Acetylated α-tubulin was used as a reference marker for axonemal microtubules, as acetylation of the luminal lysine 40 residue is a characteristic modification of stable ciliary microtubules.

U-ExM imaging and quantitative fluorescence analyses revealed distinct patterns of tubulin PTM retention on intracellular c-rings. Detyrosinated α-tubulin signals were markedly reduced, with little detectable signal remaining on c-rings at 30 minutes after partial deciliation (**Fig. 2a; Extended Data Fig. 1a, b**). Glutamylation and polyglutamylation signals were also reduced, although to a lesser extent (**Fig. 2b, c; Extended Data Fig. 1a, b**). In contrast, polyglycylation remained detectable on c-rings at levels comparable to those of acetylated α-tubulin (**Fig. 2d; Extended Data Fig. 1a, b**). These observations indicate that intracellular c-rings retain a subset of tubulin modifications characteristic of mature axonemal microtubules, despite changes in their overall modification profile. The differential retention of these modifications may influence the susceptibility of c-ring microtubules to cytoplasmic disassembly machinery. However, whether these PTM changes directly regulate axonemal breakdown remains to be determined. Together, these findings suggest that c-rings preserve features of mature ciliary microtubules during intracellular processing, raising the possibility that their clearance involves mechanisms beyond conventional microtubule disassembly.

### Proteomic profiling identifies candidate pathways associated with ciliary recovery

To explore the cellular pathways associated with c-ring processing and ciliary regeneration, we performed quantitative whole-cell proteomics on ciliated and partially deciliated *Tetrahymena*. We selected the 120-minute post-deciliation time point, when c-ring abundance had begun to decline and regenerating cilia became detectable (**Fig. 1e–g**). We reasoned that profiling protein abundance during this period could provide an overview of the cellular response to partial deciliation and identify candidate pathways potentially associated with axonemal processing and ciliogenesis. Analysis of two independently acquired tandem mass tag (TMT) datasets identified proteins with increased abundance following partial deciliation, including a subset detected reproducibly in both experiments (**Fig. 3a–c**).

**Figure 1.**
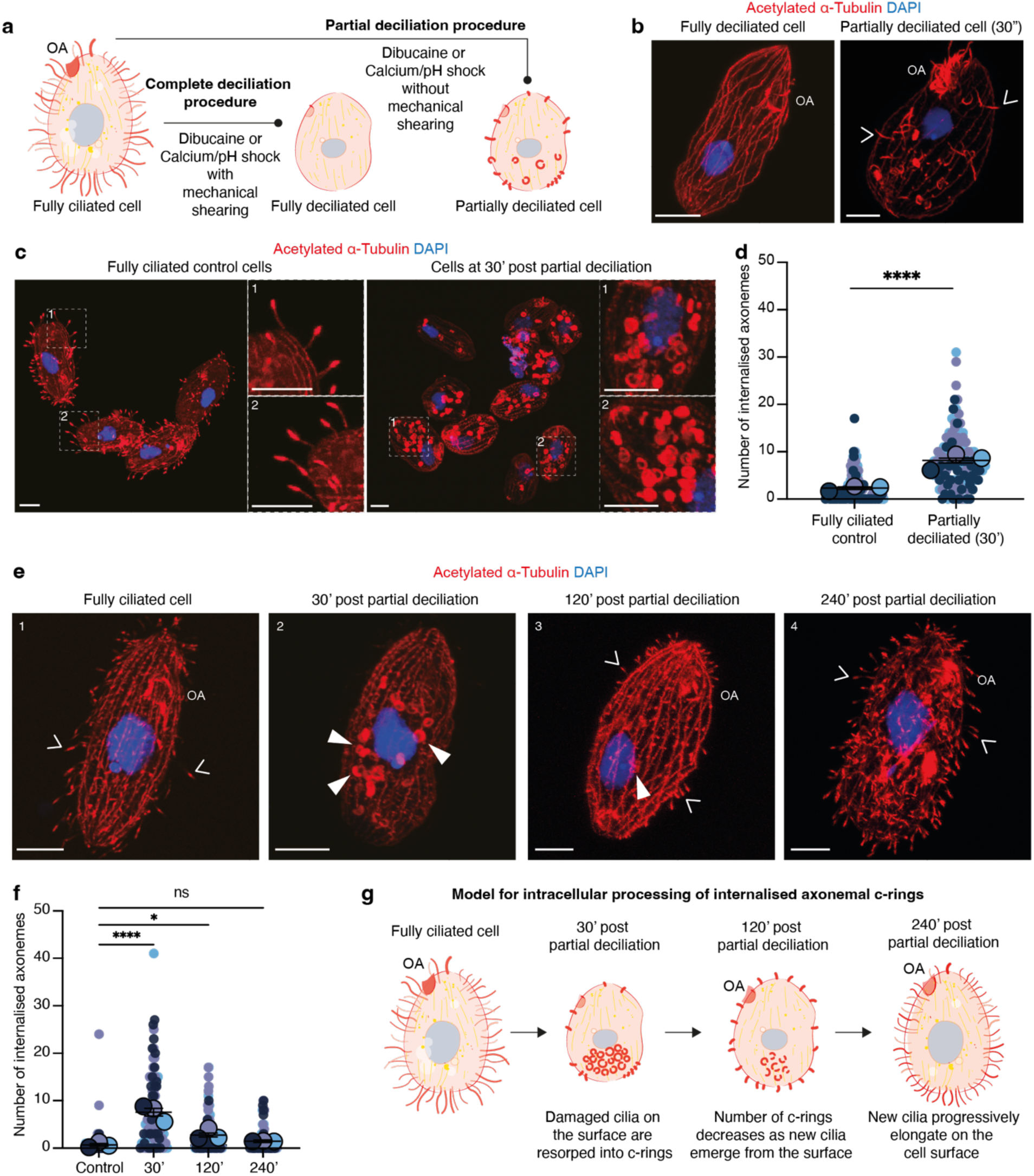
Damaged motile axonemes accumulate as intracellular rings during ciliary regeneration. **a.** Schematic showing deciliation procedures in *Tetrahymena*. Fully ciliated cells can be completely or partially deciliated by dibucaine hydrochloride or calcium/pH shock with or without mechanical shearing respectively. **b.** Representative fully and partially deciliated cells 30 seconds after calcium/pH shock, stained with acetylated α-tubulin marking ciliary axonemes (red) and DAPI marking nuclei (blue). **c.** Deciliation induces ciliary internalization in wild-type *Tetrahymena*. Fully ciliated control cells and cells 30 minutes after partial deciliation stained as in (b). **d.** C-ring numbers in control and partially deciliated cells from three replicates (n=90 cells per condition; 30 per replicate; ****p < 0.0001) shown as dot plots; bars = mean ± s.d. **e.** Time-course immunostaining of wild-type cells before and 30, 120, and 240 minutes (panels 1-4 respectively) after partial deciliation. Open and closed arrowheads indicate external cilia and internalised c-rings respectively. **f.** C-ring numbers in ciliated and partially deciliated cells at the indicated time points from three replicates (n=90 cells per time point; 30 per replicate; adjusted ****p < 0.0001, *p < 0.0127; ns = 0.6370, not significant) shown as dot plots; bars = mean ± s.d. **g.** Putative model of ciliary internalization and clearance over time. OA refers to oral apparatus. Scale bars = 10 μm (b, c, e).

**Figure 2.**
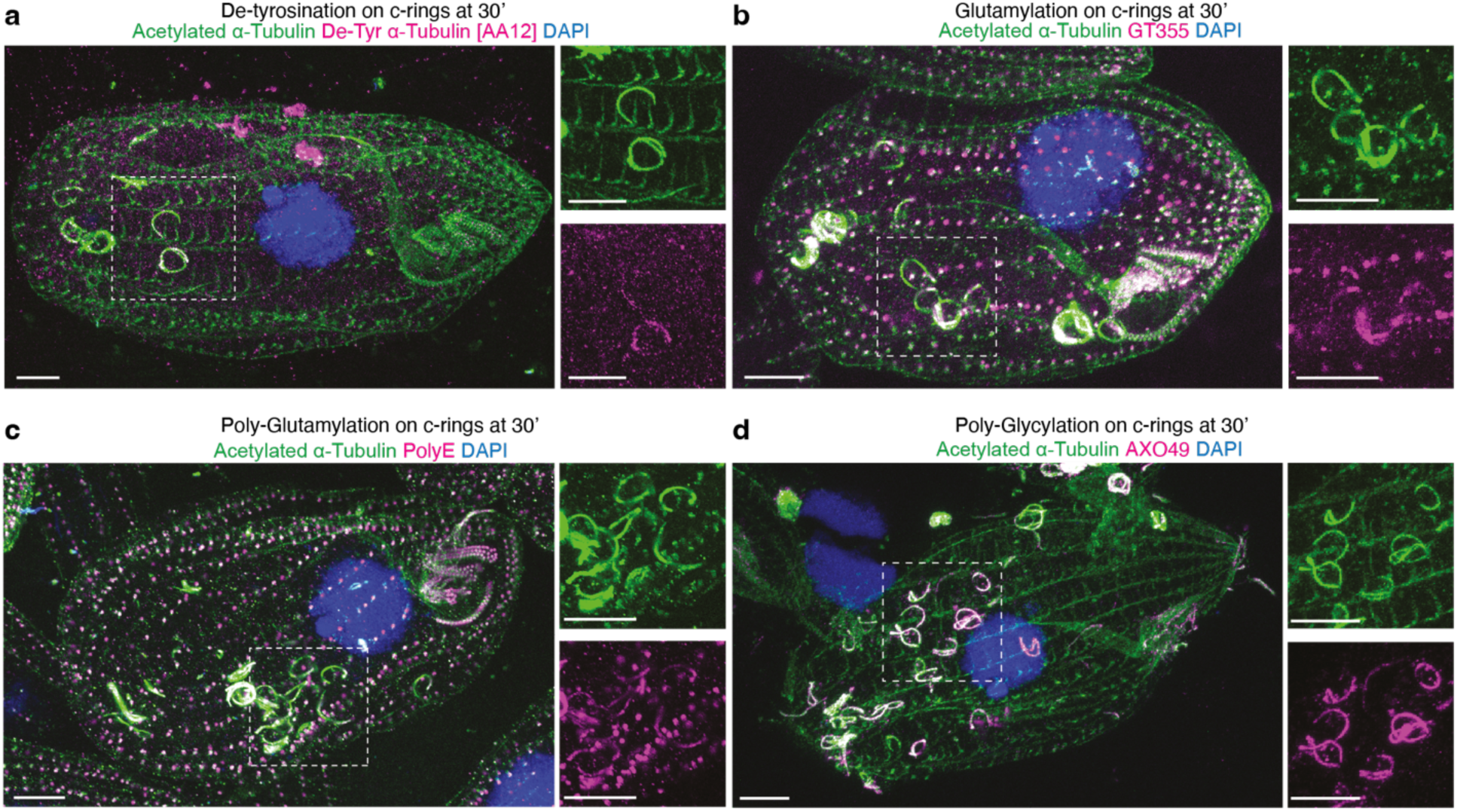
Ciliary c-rings are marked by a tubulin code characteristic of highly stable microtubule structures. **a-d.** Ultrastructure expansion microscopy (4.5x expansion) of *Tetrahymena* cells processed 30 minutes after CaCl₂-induced partial deciliation, co-immunostained for acetylated ɑ-tubulin and individual tubulin PTMs: (**a**) de-tyrosination (AA12), (**b**) glutamylation (GT335), (**c**) poly-glutamylation (PolyE), and (**d**) poly-glycylation (AXO49). Scale bar = 20 μm. Magnified views of c-rings within dotted boxes are shown on the right for each expanded cell.

**Figure 3.**
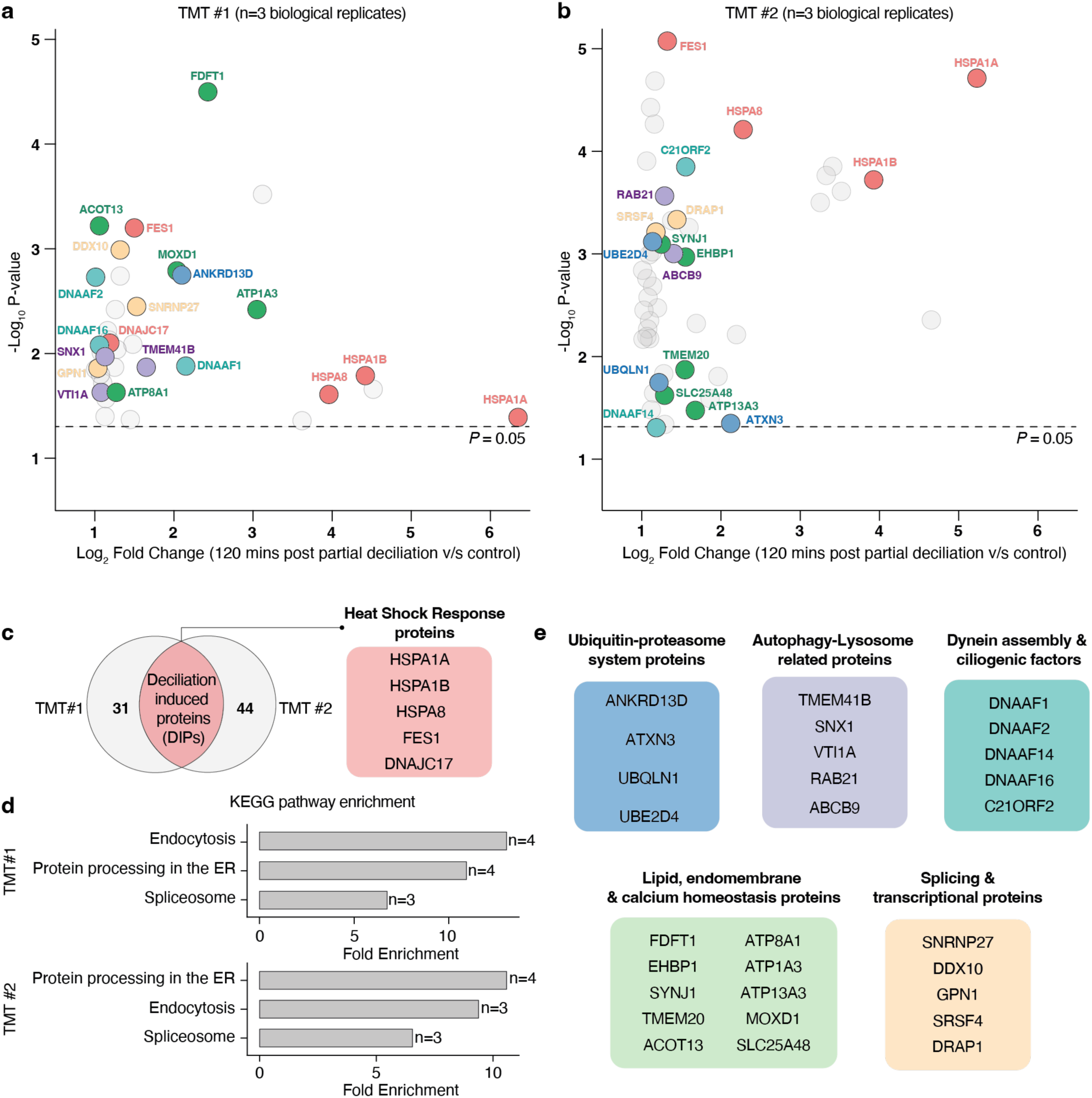
Partial deciliation is associated with changes in proteostatic and membrane-trafficking pathways. **a. b.** Whole-cell quantitative proteomic analyses of two independent TMT datasets. Scatter plots show protein enrichment 120 minutes after CaCl₂-induced partial deciliation versus ciliated controls (x-axis) and significance (y-axis; n=3 biological replicates per condition in each independent dataset). *P* values were calculated using two-tailed equal-variance *t*-tests; nominal *p* threshold = 0.05 (−log₁₀=1.3; dotted line). Upregulated proteins are colored according to categories shown in (c) and (e). Upregulated proteins not belonging to specific categories are grayed out. Proteins falling under the threshold are not shown. **c.** The Venn diagram highlights uniquely upregulated proteins and commonly upregulated deciliation-induced proteins (DIPs) which include members of the heat shock 70 protein family. **d.** Pathway enrichment analysis using ShinyGO, showing upregulation of common pathways from both independent datasets with a 0.05 FDR cutoff. Gene counts for each pathway are shown. **e.** Key upregulated proteins from (a) and (b) are grouped by functional category.

Among the proteins showing increased abundance across both datasets were several members of the HSP70 chaperone family, including proteins annotated as homologues of HSPA1A, HSPA1B and HSPA8, together with the HSP70-associated nucleotide exchange factor FES1 and co-chaperone DNAJC17 (**Fig. 3c**). Previous studies identified a group of deciliation-induced polypeptides (DIPs), including a cluster of stress-associated proteins termed sp73–75, whose synthesis was induced following deciliation and heat shock ^23^. The identification of HSP70-family proteins in our datasets is consistent with these earlier observations and suggests that partial deciliation engages a protein quality-control response with similarities to the heat-shock response.

Previous studies have shown that deciliation increases protein turnover and stimulates the synthesis of ciliary proteins, while regeneration in starved *Tetrahymena* does not require substantial de novo membrane lipid synthesis ^24^. Pathway enrichment analyses identified endocytic, proteostatic and splicing-related pathways among those over-represented in the two datasets (**Fig. 3d**). Functional annotation of proteins with increased abundance further highlighted factors associated with ubiquitin signalling, autophagy and lysosomal pathways, dynein assembly, lipid, membrane and calcium homeostasis, endomembrane trafficking, transcription and RNA processing (**Fig. 3e**). Among these were proteins annotated as homologues of TMEM41B, EHBP1, RAB21 and ANKRD13D, which have established associations with lipid homeostasis, endosomal trafficking or autophagy-related processes in other eukaryotic systems and proteins associated with lysosomal and ubiquitin-dependent protein turnover, including ABCB9, ATXN3, UBQLN1 and UBE2D4 ^25–33^. The upregulation of lipid scramblases points to remodelling of lipid membranes, evidencing the suggestion that internal membranes may be repurposed to facilitate cilia regeneration. Although these abundance changes do not establish a direct role for individual proteins in c-ring clearance, they highlight membrane trafficking and intracellular protein turnover as candidate processes that may contribute to the response to partial deciliation.

In parallel, we observed increased abundance of several proteins associated with ciliary assembly, including axonemal dynein assembly factors (DNAAFs) 1, 2, 14 and 16 (**Fig. 3e**). These factors are involved in the cytoplasmic assembly of axonemal dynein complexes and may contribute to the replenishment of ciliary components during regeneration ^34^. We also detected increased abundance of a C21ORF2 homologue, a conserved protein implicated in primary ciliogenesis and associated with NEK1-dependent pathways in other systems ^35,36^. Together, these analyses identify a broad proteomic response to partial deciliation, encompassing pathways associated with protein quality control, endomembrane trafficking and ciliary assembly. Although the contributions of individual proteins to c-ring processing remain unresolved, the enrichment of factors linked to intracellular trafficking and degradation suggested that membrane-associated pathways might participate in handling damaged axonemal material. Therefore, we next examined the ultrastructural organization of intracellular c-rings and their association with membrane-bound compartments.

### Damaged axonemes accumulate within VPS13A-positive membrane compartments

Our proteomic analyses identified changes in proteins associated with endomembrane trafficking and intracellular degradation following partial deciliation, raising the possibility that membrane-associated pathways contribute to c-ring processing. To investigate the ultrastructural organization of intracellular c-rings, we examined partially deciliated *Tetrahymena* cells using transmission electron microscopy (TEM). Compared with ciliated controls, partially deciliated cells contained large intracellular membrane-bound compartments with multilamellar features enclosing recognizable axonemal structures (**Fig. 4a; Extended Data Fig. 1a, b**). Longitudinal and transverse sections revealed axonemal profiles, including characteristic 9+2 microtubule arrangements, indicating that substantial features of motile axonemal architecture are preserved within these compartments (**Fig. 4a; Extended Data Fig. 1a, b**).

**Figure 4.**
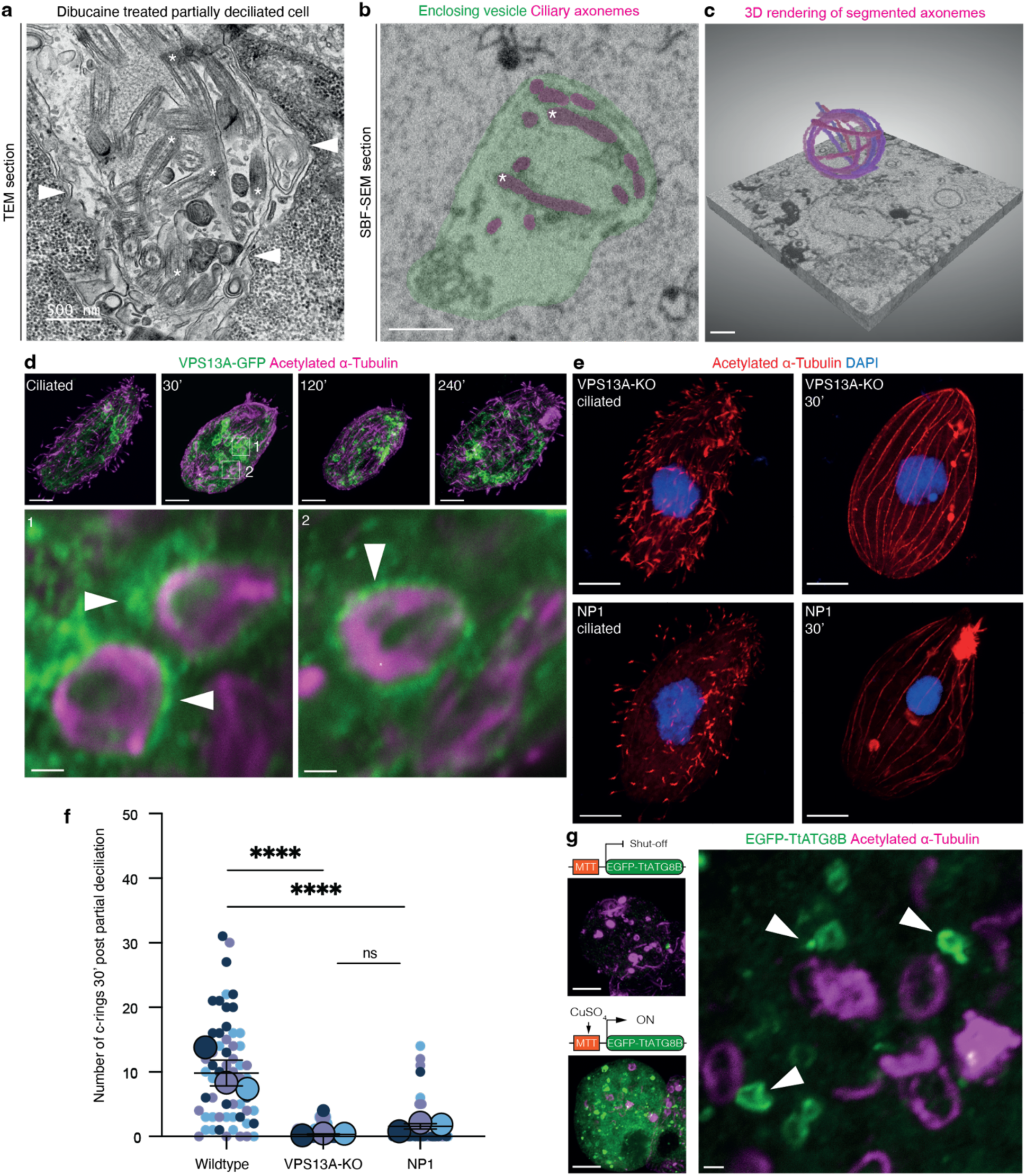
VPS13A-positive membrane compartments sequester damaged cilia. **a.** Representative transmission electron micrograph of a partially deciliated *Tetrahymena thermophila* cell showing membrane-bound compartments containing axonemal structures. Arrowheads indicate the enclosing membrane and asterisks mark axonemal profiles. Scale bar = 500 nm **b.** Representative serial block-face scanning electron microscopy (SBF-SEM) section showing an enclosing vesicle (green) containing coiled ciliary axonemes (c-rings, purple; asterisks). **c.** Three-dimensional reconstruction of the compartment shown in (b), illustrating the spatial organization of coiled axonemes within the enclosing membrane. Scale bars in (b, c) = 1 μm. **d.** Representative fluorescence images of cells expressing VPS13A-GFP (green) stained for acetylated α-tubulin (magenta), showing VPS13A-positive compartments associated with intracellular c-rings following partial deciliation. Enlarged views of the indicated regions are shown. Scale bars= 10 μm, 1 μm (bottom panels). **e**. Representative immunofluorescence images of VPS13A-KO and NP1 cells before and 30 min after partial deciliation, stained for acetylated α-tubulin. Scale bars = 10 μm. **f.** Quantification of intracellular c-ring numbers in wild-type, VPS13A-KO and NP1 cells 30 minutes after partial deciliation. Data were collected from three independent experimental replicates (n = 60 cells per condition; 20 cells per replicate). Individual cell measurements and replicate summaries are shown. P values were calculated using unpaired t-test with Welch’s correction ****p<0.0001, ns = not significant. **g**. Representative fluorescence images of cells expressing EGFP-TtATG8B (green), stained for acetylated α-tubulin (magenta). Arrowheads indicate TtATG8B-positive vesicular structures proximal to intracellular c-rings in the enlarged panel on the right. Scale bars = 10 µm (left panels), 1 µm (right panel).

To examine the three-dimensional organization of these structures, we performed serial block-face scanning electron microscopy (SBF-SEM) and volumetric reconstruction of cells 30 minutes after partial deciliation. These analyses revealed large membrane-enclosed compartments containing multiple coiled axonemal structures (**Fig. 4b, c; Supplementary Video S1**). Individual axonemes were frequently positioned along the periphery of the enclosing compartment, suggesting that spatial confinement may contribute to their characteristic coiled morphology. Together, these observations establish that intracellular c-rings can accumulate as structurally recognizable axonemal assemblies within membrane-bound compartments.

The presence of multiple axonemes within these compartments raised the possibility that phagocytic processes contribute to their intracellular sequestration. To investigate this, we examined a *Tetrahymena* strain expressing endogenously tagged VPS13A–GFP. VPS13A is a lipid-transfer protein previously shown to localise to phagosomal membranes and contribute to efficient phagocytosis in *Tetrahymena* ^37^. Following partial deciliation, we observed intracellular c-rings associated with VPS13A– GFP-positive compartments at the 30-minute time point (**Fig. 4d**). These observations are consistent with the involvement of phagosome-associated membranes in the sequestration of axonemal material.

To further investigate the relationship between phagocytic processes and c-ring accumulation, we examined two *Tetrahymena* mutants with defects in phagocytosis-associated functions. VPS13A-deficient cells exhibit impaired phagosome formation and maturation, whereas the temperature-sensitive NP1 mutant develops a disorganized oral apparatus that compromises phagocytic function^38^. Following partial deciliation, both VPS13A-deficient and NP1 cells exhibited a marked reduction in detectable intracellular c-rings compared with wild-type cells (**Fig. 4e, f**). These findings provide genetic evidence linking phagocytosis-associated processes to c-ring accumulation.

We next asked whether autophagy-associated machinery might also contribute to the processing of these membrane-enclosed axonemal structures. Previous studies have implicated VPS13-family proteins in autophagy-related membrane dynamics ^39–41^, while our proteomic analyses identified changes in proteins associated with autophagic and endolysosomal pathways. We therefore examined a *Tetrahymena* strain expressing EGFP–TtATG8B under the control of a copper-inducible promoter (**Fig. 4g; Extended Data Fig. 1a, b**). TtATG8B is an ATG8-family protein implicated in autophagy-associated processes, including mitochondrial and nuclear degradation in *Tetrahymena* ^42–44^. Following partial deciliation, we observed an increase in EGFP–TtATG8B signal and the appearance of TtATG8B-positive vesicular structures in close proximity to intracellular c-rings (**Fig. 4g; Extended Data Fig. 1a, b**). Control experiments confirmed that reporter expression depended on copper induction (**Extended Data Fig. 1a, b**). These observations suggest that autophagy-associated membranes may participate in the cellular response to intracellular axonemal accumulation, although their direct involvement in c-ring processing remains to be established.

Together, the ultrastructural, localization and genetic observations identify membrane sequestration as a prominent feature of intracellular c-ring accumulation and implicate phagocytosis-associated processes in this phenomenon. The proximity of TtATG8B-positive vesicles further raises the possibility of cooperation with autophagy-associated machinery. These findings prompted us to investigate whether intracellular c-ring accumulation is required for the subsequent regeneration of motile cilia.

### Ciliary regeneration does not require c-ring accumulation

The progressive disappearance of intracellular c-rings coincided with the re-emergence of surface cilia, raising the question of whether the processing of damaged axonemal material contributes to ciliary regeneration. The marked reduction in c-ring accumulation in VPS13A-deficient cells (**Fig. 4e, f**) provided an opportunity to investigate whether these processes are functionally linked. Previous studies have established that *Tetrahymena* can regenerate cilia using pre-existing cytoplasmic precursor pools, even when new protein synthesis is inhibited ^13^. We therefore compared ciliary regeneration in wild-type and VPS13A-deficient cells following partial deciliation, both in the presence and absence of cycloheximide (CHX), to examine whether c-ring accumulation is required for regeneration when protein synthesis is inhibited (**Fig. 5a, b**).

**Figure 5.**
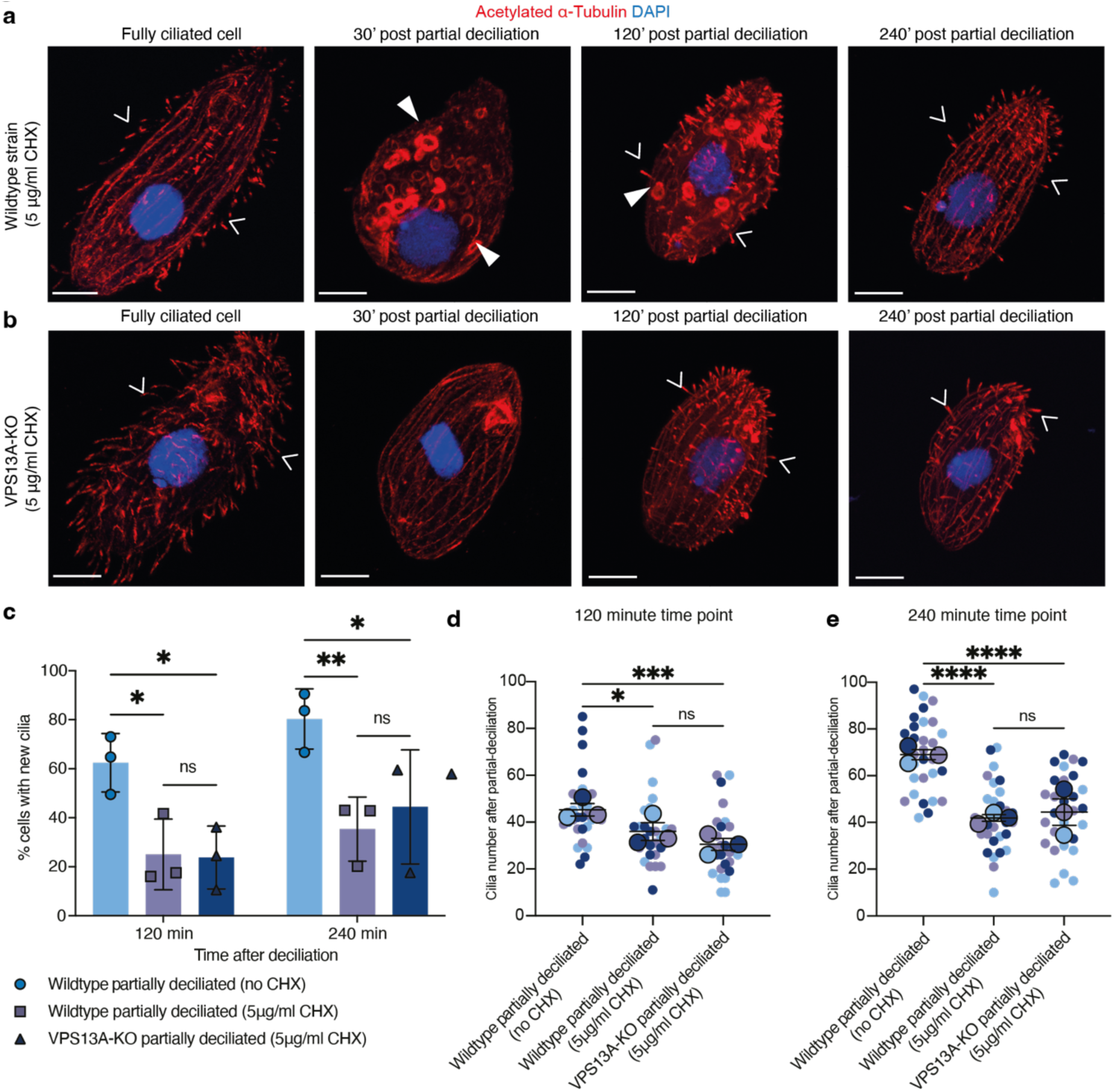
Ciliary regeneration proceeds without detectable c-ring accumulation. **a. b.** Representative immunostaining images of wildtype and VPS13A-knockout (VPS13A-KO) cells treated with 5µg/ml cycloheximide (CHX), before and at 30, 120, and 240 minutes after partial deciliation. Cells are stained with acetylated-tubulin (red) and DNA (blue). White arrows indicate external cilia. Internalised c-rings are marked with white arrowheads. Scale bar = 10 µm. **c.** Graph showing percentage of cells with new regenerated cilia at 120 and 240 minutes post-partial deciliation grown without or in the presence of 5µg/ml CHX. Independent biological replicates (n = 3) are shown, and bars show means ± s.d. For 120 min adjusted \**p* < 0.0267, \**p* < 0.0222, ns = 0.9942, not significant; for 240 min adjusted \*\**p* < 0.0089, \**p <* 0.0332, ns = 0.7493, not significant. **d. e.** Dot plots showing quantification of cilia visible per cell at 120 min (c) and 240 min (d) post-partial deciliation. Data are collected from 3 experimental replicates (n = 60 cells in total; 20 cells in each replicate per timepoint) and bars show means ± s.d. For 120 min adjusted \**p <* 0.0317, \*\*\**p <* 0.0002, ns = 0.3334, not significant. For 240 min adjusted \*\*\*\**p* < 0.0001, ns = 0.8900, not significant.

In the presence of CHX, approximately 20% of partially deciliated wild-type and VPS13A-deficient cells exhibited regenerated cilia at 120 minutes, increasing to approximately 40% by 240 minutes (**Fig. 5c**). By comparison, approximately 60% and 80% of wild-type cells exhibited cilia at the corresponding time points in the absence of CHX. Thus, CHX treatment reduced the proportion of cells exhibiting ciliary regeneration but did not completely prevent it, consistent with previous observations ^13^. Notably, we did not detect a significant difference in the proportion of cells exhibiting regenerated cilia between wild-type and VPS13A-deficient cells under CHX treatment at either time point.

To assess the extent of regeneration in individual cells, we next quantified the number of cilia in cells that had initiated ciliogenesis at 120 and 240 minutes after partial deciliation. As expected, cells recovering in the absence of CHX exhibited greater numbers of cilia than those treated with CHX (F**ig. 5d, e**). However, among cells exhibiting regenerated cilia under CHX treatment, we observed no significant difference in cilia number between wild-type and VPS13A-deficient cells at either time point.

Together, these findings indicate that detectable c-ring accumulation is not essential for at least partial ciliary regeneration under the conditions examined. The ability of VPS13A-deficient cells to regenerate cilia in the presence of CHX is consistent with a contribution from pre-existing cytoplasmic ciliary precursor pools. Although these experiments do not exclude the possibility that components from pre-existing cilia are reutilised through pathways independent of c-ring accumulation, they strongly suggest that intracellular sequestration of axonemal material and the initiation of ciliary regeneration are not coupled.

### Ciliary structural proteins persist on intracellular c-rings and associate with acidic compartments

Our observations indicate that intracellular c-ring accumulation is not required for the initiation of ciliary regeneration. However, the fate of axonemal proteins associated with these structures remained unclear. To investigate whether c-rings retain major ciliary protein complexes during intracellular processing, we first examined the distribution of outer dynein arms (ODAs) using a previously validated antibody recognizing the ODA holoenzyme ^45^. ODA immunoreactivity was detected on both surface cilia and intracellular c-rings throughout the recovery time course (**Fig. 6a**). We also noted few ODA puncta separate from c-rings at the 30-minute timepoint representing early foci of new dynein synthesis and assembly and consistent with an upregulation in DNAAFs ^46^. Quantification of ODA fluorescence relative to acetylated α-tubulin showed that ODA-associated signals persisted on c-rings during the period of intracellular accumulation (**Fig. 6b**). Overall, these observations indicate that c-rings retain components of the axonemal motor machinery following partial deciliation, consistent with the preservation of recognizable axonemal architecture observed by electron microscopy.

**Figure 6.**
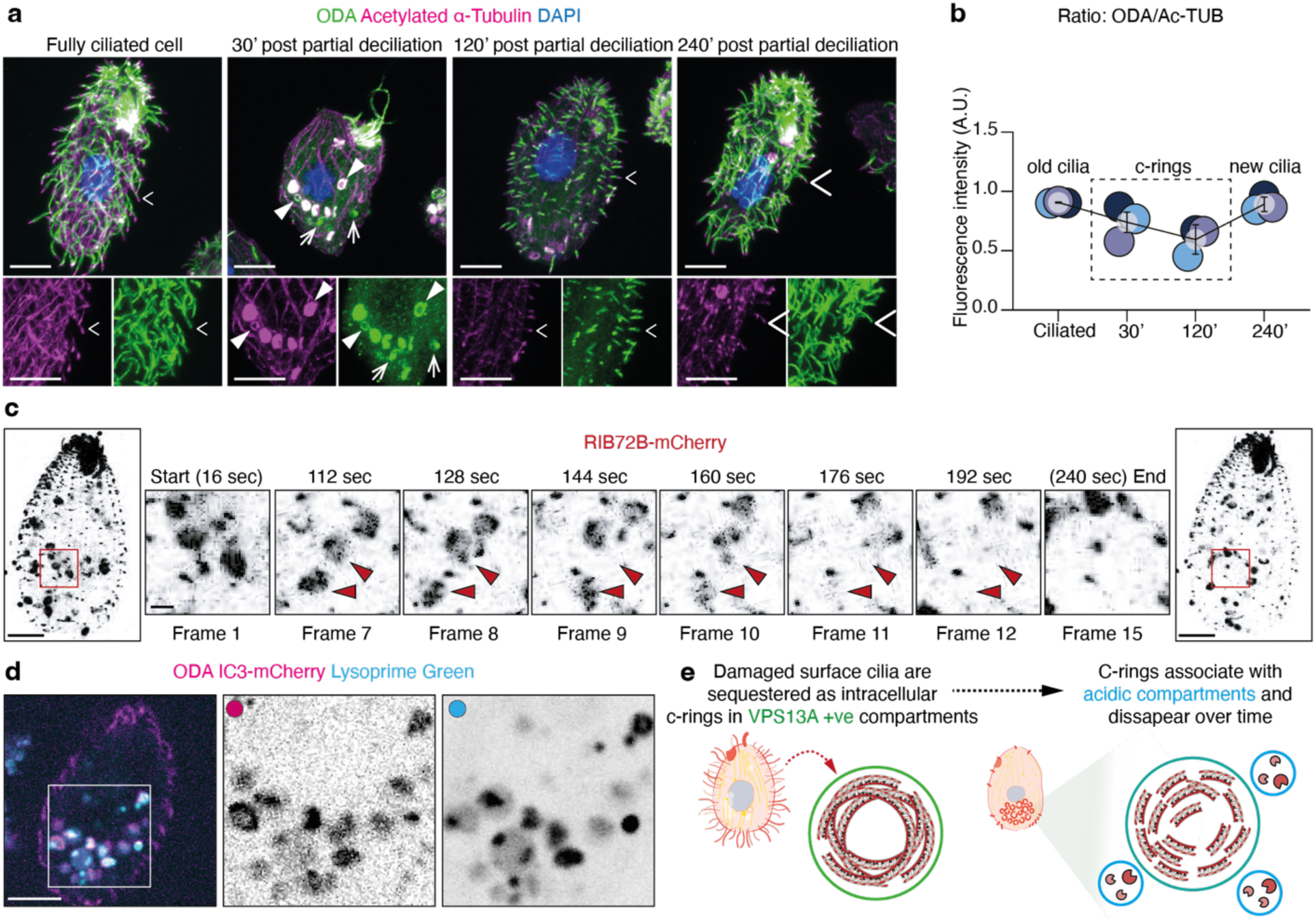
Structural ciliary proteins remain bound to c-rings and co-localise with cytoplasmic acidic compartments over time. **a.** Time-course immunostaining for wild-type *Tetrahymena*. Ciliated and partially deciliated cells at 30-, 120-, and 240-minutes (panels 2-4 respectively) after CaCl_2_ treatment, were stained with ODA (green) and acetylated ɑ-tubulin (magenta) antibodies. White open arrowheads indicate external cilia, internalised c-rings are marked with white closed arrowheads, and white arrows point to puncta. Scale bar = 10 µm. **b.** Quantification of ODA fluorescence intensity signals on external cilia and internal c-rings (dashed box) relative to acetylated ɑ-tubulin shown as a ratio. **c.** Individual time frames from live-cell time-lapse Movie S2 of a *Tetrahymena* cell strain expressing RIB72B-mCherry are shown with the time stamps indicated. Red boxes and red arrowheads are used to highlight the gradual loss of RIB72B-mCherry signal from c-rings over time. Scale bars = 10 µm (enlarged panels) and 2 µm (zoomed panels). **d.** Representative still image of ODA IC3-mCherry co-localising with LysoPrime green from a cell that has regained new cilia and motility (Supplementary Video S3). Higher magnification views of the boxed regions for each channel are shown. **e.** Proposed model for the sequestration of internalised damaged cilia and routing to acidic compartments for clearance.

To further examine the behaviour of c-ring-associated proteins, we performed live-cell imaging using a strain expressing mCherry-tagged RIB72B, a microtubule inner protein associated with the A-tubule of axonemal doublet microtubules ^47^. RIB72B-mCherry-labelled intracellular structures resembling c-rings were observed following partial deciliation (**Fig. 6c; Supplementary Video S2**). During time-lapse imaging, the fluorescence signal associated with individual structures progressively diminished over time. To explore the possible association of ciliary material with acidic compartments, we imaged IC3 mCherry-expressing cells labelled with LysoPrime Green. IC3 is an essential subunit of the ODA complex, and this strain was used to track ODA distribution during c-ring clearance. We observed instances of close spatial association between IC3-mCherry-positive structures and LysoPrime-labelled compartments (**Fig. 6d; Supplementary Video S3**). Together with the ultrastructural and VPS13A localization data, these findings support a model in which intracellular c-rings are sequestered within membrane-bound compartments and subsequently associate with acidic compartments, consistent with their processing through a phagolysosomal clearance pathway (**Fig. 6e**).

## Discussion

The fate of damaged motile cilia presents a fundamental challenge for cellular proteostasis. These organelles comprise highly stable microtubule-based assemblies containing hundreds of structural and regulatory proteins yet must be remodeled or replaced when their function is compromised. Here, we identify an intracellular route through which damaged motile axonemes are sequestered within membrane-bound compartments and directed towards degradation during ciliary regeneration. Following acute deciliation in *Tetrahymena thermophila*, axonemal structures accumulate as intracellular coils, which we term c-rings, before progressively disappearing as new cilia emerge. Ultrastructural imaging reveals that these coils are enclosed within large intracellular vesicles, while their association with phagocytic machinery and acidic compartments supports a pathway for their degradative processing. These findings reveal an unexpected fate for damaged motile axonemes and establish intracellular sequestration as a mechanism for managing the turnover of large, stable cytoskeletal assemblies.

A notable feature of this pathway is the preservation of axonemal architecture during the early stages of intracellular processing. C-rings retain structural components, including outer dynein arms and the microtubule inner protein RIB72B, together with a subset of tubulin post-translational modifications characteristic of mature ciliary microtubules. Their persistence suggests that damaged axonemes are not immediately dismantled into soluble subunits before intracellular sequestration. Instead, the enclosure of multiple axonemal structures within membrane-bound compartments provides a potential mechanism for isolating complex cytoskeletal cargo from the surrounding cytoplasm. The accumulation of c-rings in VPS13A-positive compartments, together with their marked reduction in phagocytosis-defective mutants, implicates phagocytic trafficking in this process. The subsequent association of ciliary proteins with acidic compartments is consistent with delivery to a degradative pathway. Although the precise sequence of membrane remodelling and cargo breakdown remains to be established, these observations suggest that bulk compartmentalisation can provide an alternative to the complete cytosolic disassembly of stable axonemal structures.

Our findings also raise the possibility of cooperation between phagocytic and autophagic machinery during damaged axoneme clearance. Quantitative proteomics identifies a coordinated response involving protein quality control, endomembrane trafficking and autophagy-associated factors following deciliation. Moreover, TtATG8B-positive vesicles accumulate near internalised c-rings, suggesting that autophagy-related membranes may participate in their processing. Such cooperation would be consistent with emerging evidence that phagosomes and autophagosomes can interact during the degradation of unusually large or complex cargo ^48^. Nevertheless, the proximity of TtATG8B-positive structures does not establish membrane fusion or a requirement for canonical autophagy. Whether these compartments undergo direct fusion, recruit autophagy-associated proteins to phagosomal membranes, or represent parallel degradative pathways remain important questions. Resolving these alternatives will help define how cells adapt established membrane-trafficking systems to the disposal of large cytoskeletal organelles.

An important implication of our study is that the intracellular accumulation of damaged axonemes is not required for ciliary regeneration. Although c-ring disappearance coincides with the re-emergence of surface cilia, cells defective in c-ring accumulation retain the capacity to regenerate cilia, including when protein synthesis is inhibited by cycloheximide. These observations support the existence of a pre-existing cytoplasmic precursor pool capable of sustaining at least partial ciliogenesis independently of detectable c-ring sequestration. This finding distinguishes the disposal of damaged axonemal structures from the initiation of replacement cilium assembly. However, our experiments do not exclude the possibility that some ciliary proteins are recycled through pathways that do not involve detectable c-rings, nor do they establish the quantitative contribution of salvaged proteins to regeneration under physiological conditions. Direct tracking of pre-existing axonemal proteins into newly assembled cilia will be required to resolve these possibilities. Nevertheless, the capacity to regenerate cilia without c-ring accumulation indicates that the two processes are not obligatorily coupled.

The intracellular processing of damaged motile axonemes may represent a broader principle of organelle quality control. Previous studies have identified autophagy-associated turnover of ciliary components in mammalian cells, while intracellular accumulations of axonemal structures have been observed in respiratory epithelia following viral infection ^8–10^. Our findings extend these observations by identifying membrane-enclosed, structurally recognizable motile axonemes and linking their intracellular fate to phagocytic and acidic compartments. Whether comparable pathways operate in mammalian multiciliated epithelia, and whether they contribute to recovery from infection, inflammation or mechanical injury, remain open questions. More broadly, our study suggests that cells can respond to organelle damage through two distinguishable activities: sequestration and disposal of compromised structures, and mobilization of precursor material for organelle replacement. This separation provides a framework for understanding how cells maintain the integrity of complex motile organelles during recovery from stress.

## Extended data figures and legends

**Figure S1.**
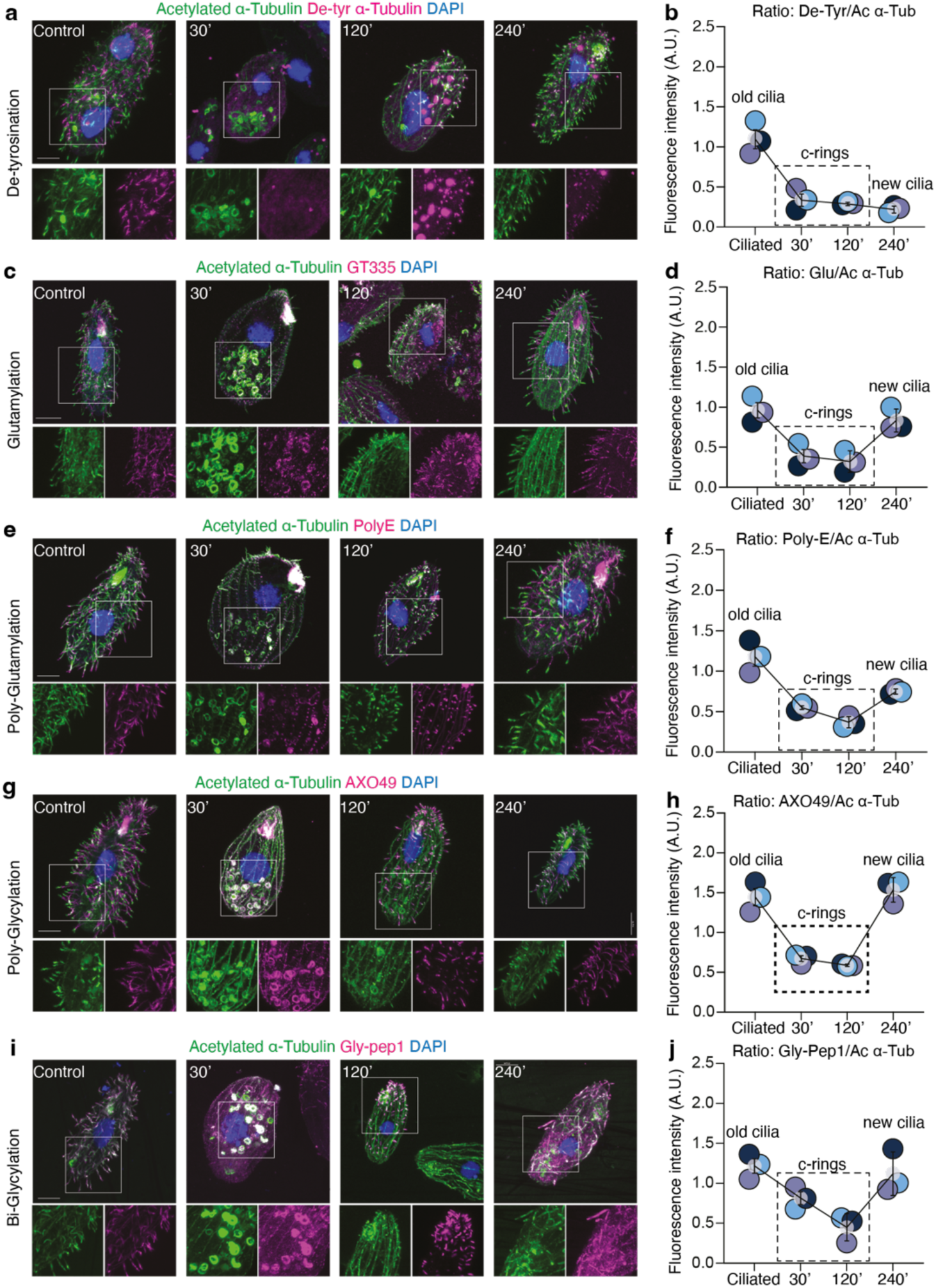
Quantification shows c-rings retain microtubule stabilising tubulin post-translational modifications. **a-j.** Representative time course immunostaining images and quantification of PTM levels on c-rings relative to acetylated ɑ-tubulin (PTM/ ɑ-tubulin ratio) measured using confocal images of ciliated cells and cells at 30-, 120- and 240-minutes after CaCl₂-induced partial deciliation. Cells were co-immunostained with antibodies for acetylated ɑ-tubulin and individual tubulin PTMs: (a, b) de-tyrosination, (c, d) glutamylation (GT335), (e, f) poly-glutamylation (PolyE), (g, h) poly-glycylation (AXO49), and (i, j) glycylation (Gly-pep1). White boxes indicate magnified views shown under each image. Scale bar = 10 μm. For the quantification graphs (b, d, f, h, j) each dot represents an average value from three independent immunostaining experiments (n = 3 experimental repeats; measurements from 10-15 axonemes per cell with 8-10 cells within each repeat). Data points from quantifications for internalised axonemes in cells at the 30-minute and 120-minute post-partial deciliation timepoint are enclosed by dashed boxes; data points from ciliated cells and cells at 240-minute post-partial deciliation timepoint correspond to old and newly formed external surface cilia respectively.

**Figure S2.**
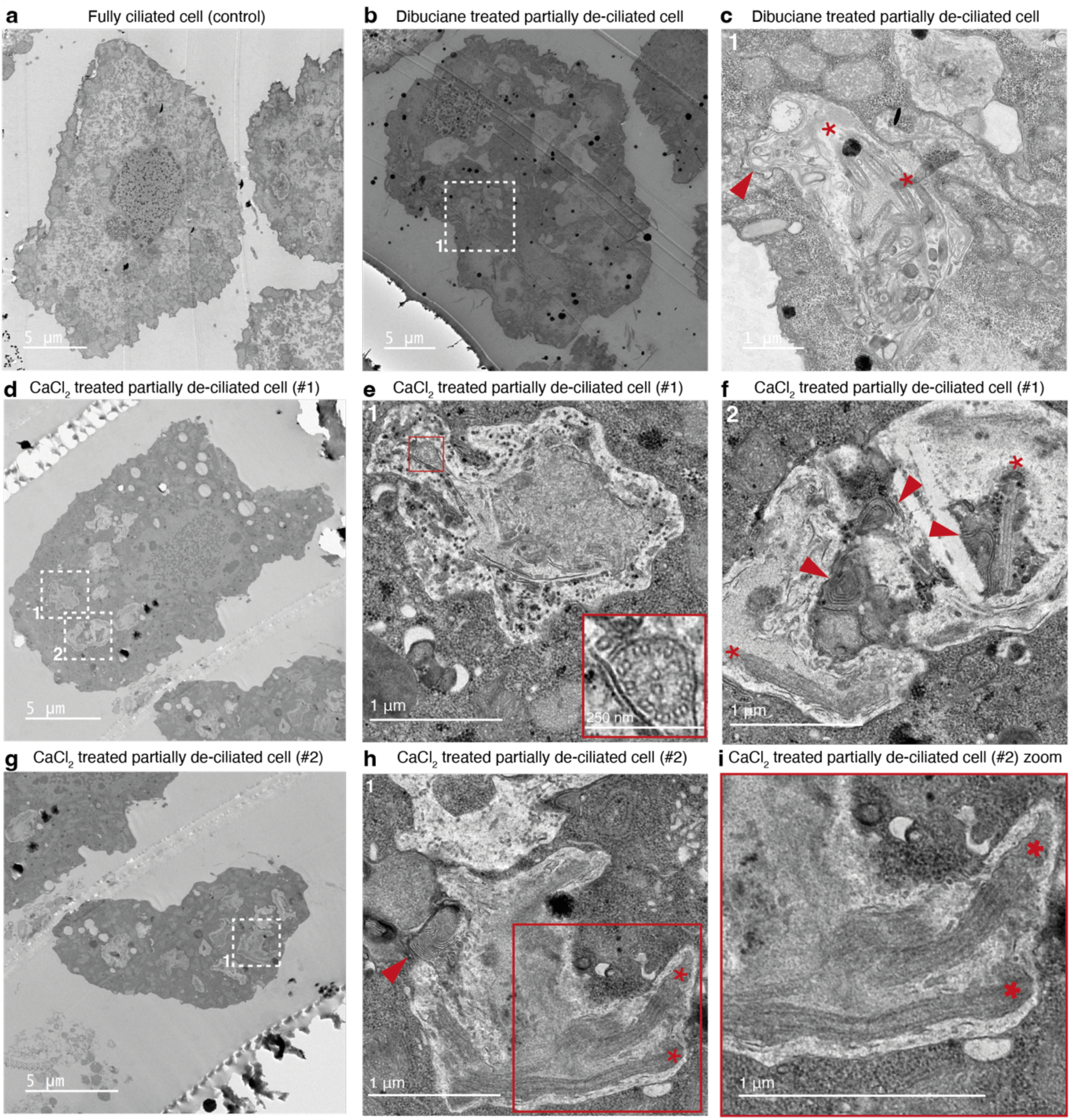
Partially deciliated cells sequester damaged axonemes in large vesicles with multilamellar features. **a. b. d. g.** Transmission electron micrographs of a ciliated *Tetrahymena* cell (a) or partially deciliated cells (b, d, g) processed 30 minutes after CaCl_2_ treatment. Scale bars = 5 μm. **c. e. f. h. i.** Higher-magnification views of regions indicated (white dashed boxes in b, d, g). Axonemal fragments (red asterisks) enclosed within cytoplasmic compartments with multilamellar features (red arrowheads) are shown. Inset in (e) is a magnified view of the boxed region showing a cross-sectional 9+2 doublet arrangement typical of motile axonemes. A magnified view of inset in (h) is shown in (i). Scale bars = 250 nm (inset in e) and 1 μm as indicated.

**Figure S3.**
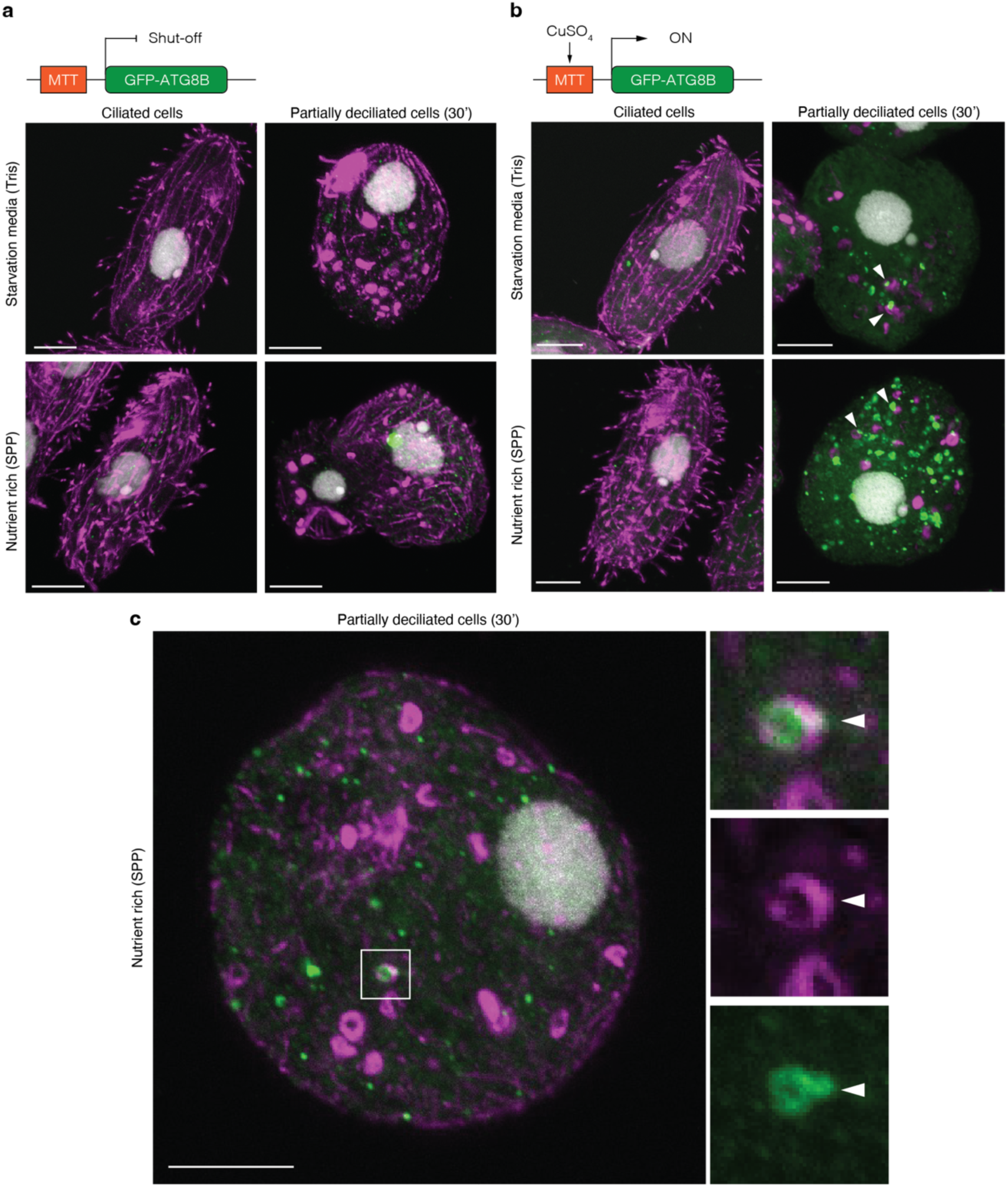
Representative immunostaining images of EGFP-TtATG8B strain grown in the presence or absence of CuSO_4_. **a. b.** Schematic of the EGFP-TtATG8B strain in which expression is controlled by a CuSO_4_ sensitive MTT2 promoter; absence of CuSO_4_ in the media shuts-off expression and its presence switches on expression. Representative immunostaining images of ciliated cells as controls and cells 30 min after partial deciliation are shown. Cells were maintained in starvation medium (10 mM Tris-HCI) or nutrient-rich SPP medium. TtATG8B-EGFP expression was shut off in the absence of CuSO_4_ (a) and induced with 0.1 mM CuSO_4_ (b). Cells immunostained with antibodies to EGFP (green), acetylated α-tubulin (magenta) and counterstained with DAPI (gray) are shown. White arrowheads in (b) indicate EGFP-TtATG8B vesicles close to c-rings. Scale bars = 10 μm. **c.** Enlarged view of a CuSO_4_-induced cell in SPP medium 30 min after partial deciliation. The boxed region is shown at higher magnification with the arrowheads pointing to a close contact between a c-ring and an EGFP-TtATG8B vesicle. Scale bars = 10 μm.

**Figure S4.**
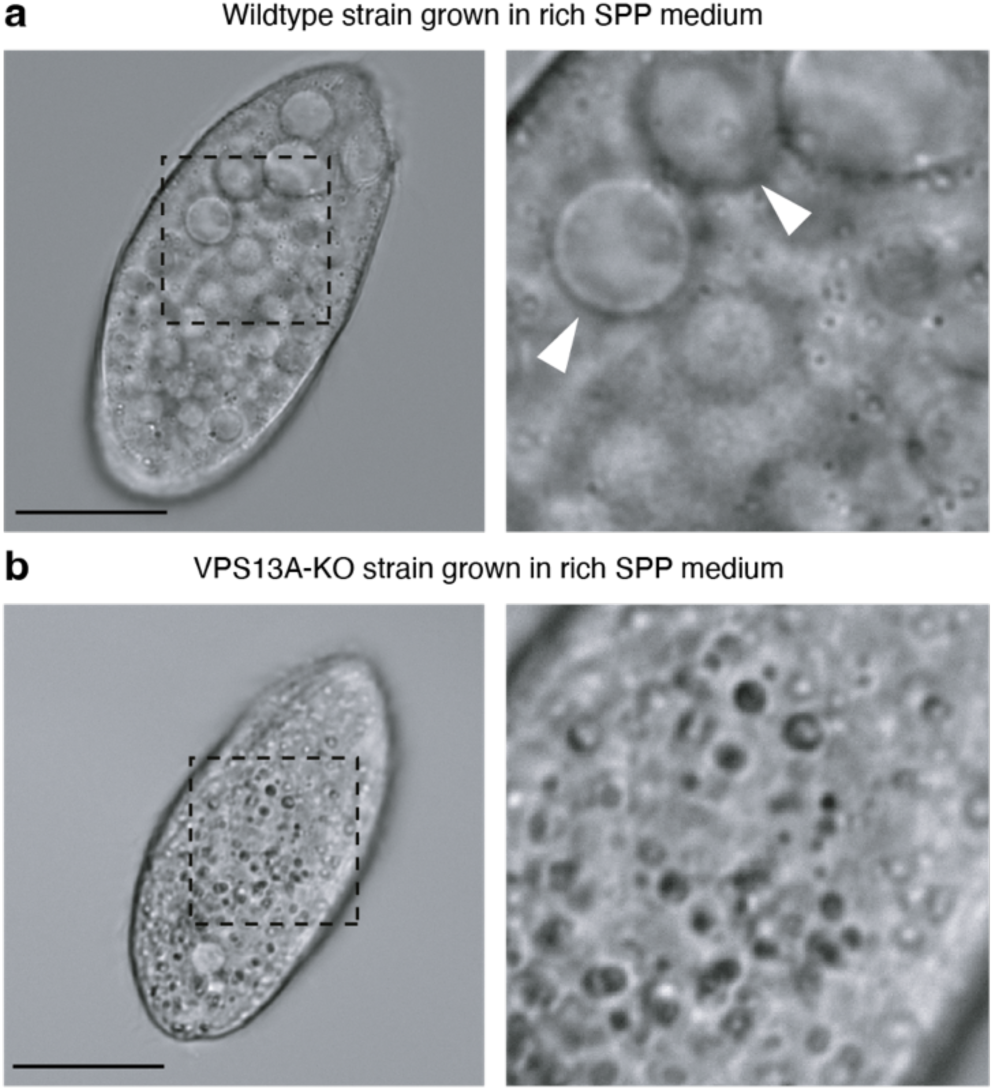
Altered food-vacuole (phagosome) morphology in VPS13A-deficient *Tetrahymena*. **a. b.** Representative brightfield images of wild-type (a) and VPS13A-KO (b) *Tetrahymena thermophila* cells cultured in nutrient-rich SPP medium. Enlarged views of the boxed regions are shown. Arrowheads in (a) indicate prominent intracellular food vacuoles observed in wild-type cells, which are markedly reduced in VPS13A-KO cells under the conditions examined consistent with the previously reported involvement of VPS13A in phagosome formation and efficient phagocytosis. Scale bars = 20 µm.

**Source Data for Figures 1, 4, 5, 6, Extended Data Figure 1:** Numerical raw data and statistical analyses used to generate graphs.

**Source Data for Figure 3:** MS raw source data for processed TMT Run #1 and Run #2 proteomic data.

**Supplementary Video S1: SBF-SEM serial sections and 3D reconstruction of axonemes.**

SBF-SEM reveals a large intracellular enclosing vesicle containing multiple c-rings. The left panel shows the three-dimensional reconstruction of the enclosing vesicle (green) and the spatial arrangement of the coiled axonemes (magenta) enclosed in the form of c-rings. The video relates to Fig. 4b, c. The right panel shows magnified serial SBF-SEM sections through the same enclosing vesicle, with individual c-rings highlighted in magenta. Scale bar = 1 μm.

**Supplementary Video S2: Live cell time-lapse imaging of RIB72B-mCherry 30 minutes post-partial deciliation.**

RIB72B-mCherry positive c-rings imaged in a live partially deciliated cell 30 minutes after calcium chloride treatment immobilised in low melting point agarose. The time-lapse sequence was captured using a SoRa spinning disk imaging system with a total length of 241 seconds and relates to frames shown in Fig. 6c. Scale bar = 20 μm.

**Supplementary Video S3: IC3-mCherry strain imaging with Lysoprime Green dye 60 minutes post-partial deciliation.**

IC3–mCherry positive c-rings (red) and LysoPrime Green-labelled acidic compartments characteristic of phagolysosomes (green) were imaged 60 minutes after calcium chloride induced partial deciliation using a SoRa spinning-disk imaging system with a total length for the time-lapse capture of 44 seconds. The yellow box tracks the selected cell shown in Fig. 6d. Scale bar = 20 μm.

## Methods

### *Tetrahymena thermophila* growth and culture conditions

All strains used in this study were obtained from the Tetrahymena Stock Centre or acquired as gifts from various labs. Wild-type (CU428; TSC_SD00178), RIB72B-mCherry (Gift from Prof. Mark Winey), VPS13A-GFP (TSC_SD01757), VPS13A-NEO (TSC_SD017158), EGFP-TtATG8B shut-off, NP1(TSC_SD01422) and IC3-mCherry strains were cultured in a modified SPP medium containing 0.25% bactoproteose peptone, 0.25% yeast extract, 0.55% glucose and 33.3 µM FeCI_3_. The cultures were maintained at room temperature without shaking. For each experiment, a high cell density, measured as OD_600_ above 1.0 was used with all cells sourced from the same flask. To induce EGFP-TtATG8B expression in the shut-off strain, cells were cultured overnight in 10 mM Tris-HCI containing 0.1mM CuSO_4_. NP1 is a temperature-sensitive oral apparatus development mutant, cells were cultured at 37°C for three to four days to induce the mutant phenotype.

### Calcium chloride based complete deciliation and partial deciliation experiments

Before each deciliation experiment, *Tetrahymena* cells were examined under a microscope to ensure they were healthy. Normal, healthy cells appear oval and swim rapidly. Cells were cultured with starvation medium (10 mM Tris-HCI, pH 7.4) overnight. Partial deciliation was achieved using the same procedure employed by Rannestad (1974). Before deciliation, cells were concentrated by centrifugation at 1100 x g for 1 minute. 1 ml of the concentrated cell suspension was added to 5.0 ml of medium A (10 mM EDTA; 50 mM sodium acetate, pH 6.0) in a 50 ml centrifuge tube and mixed by swirling. After 30 seconds, 2.5 ml cold distilled water was added, followed by the addition at 90 seconds of 0.5 ml of 0.2 M calcium chloride (CaCl_2_). After adding CaCl_2_, the suspension was mixed by inverting the tube several times. After 3 minutes and 30 seconds, the cell suspension was passed through a 10 ml syringe fitted with a large bore 19-gauge needle 2-4 times for gentle mechanical shearing to fully deciliate the cells. Partial deciliation was achieved by omitting the shearing forces at the final step. 5 minutes after full or partial deciliation, the cells were resuspended in 20 volumes of starvation medium (10 mM Tris-HCl, pH 7.4) to allow recovery and ciliary regeneration at room temperature without shaking. Phagocytosis deficient mutants were partially deciliated as above.

### Dibucaine deciliation of *Tetrahymena* cells

Concentrated cells were spun down at 1100xg for 1 minute and resuspended in half of their growth volume of starvation medium (10mM Tris-HCI, pH7.4). Dibucaine was added to achieve a final concentration of 600µM, and the cells were incubated for 4 minutes 30 seconds to 5 minutes. At the 3-minute point, cell motility could be checked under the microscope to determine if the cells were still moving or had lost motility. To stop the deciliation process, fresh starvation media was added to twice the volume immediately to dilute the dibucaine, followed by centrifugation at 1100xg for 1 minute. The supernatant was removed, and the cells were washed once with 10 mM Tris-HCI, pH 7.4. The deciliated cells were then resuspended in the original volume of fresh starvation media for the recovery period.

### Immunostaining

*Tetrahymena* cells were pelleted by centrifugation at 1100 x g for 1 minute, and the supernatant was removed. The cells were then washed by adding 2 ml of 1 x PBS and pelleted again at 1100 x g for 1 minute. After removing the supernatant, the cells were prepared for fixation by resuspending them in 4% paraformaldehyde (PFA) for 10 minutes. For centrin staining, cells were permeabilised with 0.5% Triton-X-100 in PHEM buffer and then fixed in 2% PFA in PHEM buffer. Following fixation, the cells were pelleted and submerged in 0.1% Triton X-100 (in 1x PBS) for 10 minutes to permeabilise the cell membrane. The supernatant was removed and the cells were blocked in 10% donkey serum in PBST (PBS + 0.1% Triton X-100) for 1 hour followed by antibody staining. The primary antibody was prepared in 1% donkey serum in PBST, and cells were incubated in the antibody solution for 1 hour at room temperature. The cells were washed three times with PBST for 2 minutes each. Cells were incubated with appropriate secondary antibodies diluted in 1% donkey serum in PBST for 1 hour at room temperature. Cells were then washed three times in PBST (2 minutes each). After washing, cells were gently resuspended and mounted onto glass slides using ProLong Gold antifade reagent with DAPI (Thermo Scientific) under a glass (0.16-0.19 mm) coverslip. Excess buffer was removed prior to mounting. Slides were allowed to air dry before imaging.

For immunofluorescence (IF) staining, the following primary antibodies were used at a dilution of 1:500: mouse monoclonal anti-α-tubulin (70-102, Bioacademia), rabbit polyclonal anti-acetylated α-tubulin (PA5-105102, Thermo Fisher Scientific), goat anti-acetylated α-tubulin (ab289875, Abcam), mouse anti-polyglutamylated tubulin (GT335; AG-20B-0020-C100, AdipoGen), rabbit anti-polyglutamate chains (PolyE; AG-25B-0030-C050, AdipoGen), mouse anti-detyrosinated α-tubulin (ab254154, Abcam), mouse anti-pan-polyglycylated tubulin (AXO49; MABS276, Merck), rabbit anti-glycylated tubulin (Gly-pep1; AG-25B-0034, AdipoGen), custom-generated rabbit anti-ODA (Eurogentec), and mouse anti-GFP (CSB-MA000051M1, CUSABIO). For fluorescence detection, donkey secondary antibodies conjugated to Alexa Fluor 488 (anti-mouse; A-21202, Invitrogen), Alexa Fluor 568 (anti-rabbit; A-10042, Invitrogen), and Alexa Fluor 647 (anti-goat; A-32849, Invitrogen) were used at a dilution of 1:800.

### Protein synthesis block using Cycloheximide (CHX)

*Tetrahymena* cells were starved in 10mM Tris-HCI overnight. Before the deciliation, cells were treated with 5µg/ml cycloheximide (CHX, Merck) for 30 minutes before the deciliation. After the deciliation, deciliated cells were incubated in 5µg/ml CHX for recovery.

### Confocal microscopy

Confocal images were acquired using OLYMPUS FV1000 and FV3000 Microscope, using 20x air, 60x oil, and 100x oil objectives. Microscope parameters were controlled using OLYMPUS FV software. 488 nm, 568 nm and 647 nm fluorescence channels were used for imaging, and the laser density was set at the same level before imaging for each slide. Z-stacks (∼1-2 µm optical slices) _were acquired to image the_ internalised c-rings within the cytoplasm; the depth was around 10 µm.

### Ultrastructure Expansion Microscopy (U-ExM)

U-ExM was conducted using a previously published protocol. Briefly, cells were fixed in 4% PFA on poly-D-lysine coated 12 mm coverslips. Cells were then anchored using a 1.4% formaldehyde + 2% acrylamide solution and incubated at 37°C for 3 hours. Gelation was performed using a monomer solution (19% sodium acrylate, 10% acrylamide, 0.1% N, N’-methylenebisacrylamide, TEMED and APS in PBS), followed by denaturation in a denaturation buffer (200 mM SDS, 200 mM NaCI, 50 mM Tris-HCI, pH 9) at 95°C for 1.5 hours. Gels were washed by water and expanded in water overnight. The expansion factor was calculated after expansion. Expanding gels were washed in PBS and cut into quarters for antibody incubation. Cells were incubated in primary antibodies diluted in PBS-BSA 2% for 2.5 hours at 37°C with agitation followed by a PBST (PBS + 0.1% Tween) wash. For ultrastructure expansion microscopy (U-ExM), antibodies against α-tubulin, acetylated α-tubulin (PA5-105102), polyglutamylated tubulin (GT335), polyglutamate chains (PolyE), detyrosinated α-tubulin, and pan-polyglycylated tubulin (AXO49) were used at a dilution of 1:250. Cells were incubated in appropriate secondary antibodies with DAPI at 1 mg/ml (used at 1/1000 dilution; Thermo Scientific) for 2 hours at 37°C with agitation. Donkey secondary antibodies conjugated to Alexa Fluor 488 (anti-mouse; A-21202, Invitrogen), Alexa Fluor 568 (anti-rabbit; A-10042, Invitrogen), and Alexa Fluor 647 (anti-goat; A-32849, Invitrogen) were used at a dilution of 1:800. Gels were placed in beakers with water for final expansion overnight. For imaging, gels were placed in a glass-bottom chamber and orientated. The chamber was coated in advance to avoid the gel floating on the glass.

### Live cell spinning-disk imaging of RIB72B-mCherry and IC3-mCherry strains

For immobilization of *Tetrahymena* cells, cells were briefly pelleted by low-speed centrifugation using a benchtop microfuge, and the supernatant was retained as a conditioned medium. For RIB72B-mCherry imaging, the concentrated cell pellet was gently resuspended in pre-warmed 3% low-melting-point agarose maintained at 37°C. A 50 μl aliquot of the cell–agarose mixture was immediately transferred onto a pre-warmed (37°C) glass-bottom dish and flattened using a sterile resin block to generate a thin agarose layer suitable for imaging. The gel was rapidly solidified by brief cooling on ice, after which the weight was removed, and 100 μl conditioned medium was added to overlay the gel. Imaging was initiated immediately after immobilization. Fast time-lapse movies were acquired (0.57 seconds/frame) using an Olympus SoRa super-resolution spinning-disk confocal microscope controlled by CellSens software (v4.3.1). Images were acquired using a 60× oil objective (calibrated pixel size: 183.3 nm per pixel) and an sCMOS camera. mCherry fluorescence was excited with a 561 nm OBIS laser (at 30% laser power) and detected using a 561–585 nm emission filter. The exposure time was set to 400 ms per optical section. Time-lapse imaging was performed continuously for 241 seconds. A total of 15 sequential z-stacks were collected (∼16 s per stack), each consisting of 28 optical sections with 1 µm spacing. No overt photobleaching or morphological alterations were observed during the imaging period. The IC3-mCherry strain was imaged similarly but without being resuspended in pre-warmed 3% low-melting-point agarose. A suspension of freshly deciliated IC3-mCherry cells was incubated with Lysoprime green dye (1/1000) (Dojindo, Code L261) and washed twice in 10 mM Tris-HCl. A large droplet of a suspension of deciliated cells in 10mM Tris-HCl was added to a glass bottom dish and contained within an imaging area demarcated by a lipophilic pen. Fast time-lapse movies were acquired 60 minutes after calcium chloride induced partial deciliation using a SoRa spinning-disk imaging system with a total length for the time-lapse capture of 44 seconds using the above parameters. A continuous dual color imaging setup was used. 10 z-stacks within each 44 second acquisition, followed by separate acquisitions at 15-min intervals were acquired up to 2 hours. Most cells had regained partial motility by 60 minutes but also retained c-rings that could be detected co-localising with LysoPrime green positive acidic compartments.

### Quantification of images and statistical analyses

All statistical analyses and graph generation were performed using GraphPad Prism v10. For quantification of internalised c-rings, five fields of view were randomly selected per biological replicate for image acquisition. From each field, six cells were selected for analysis, yielding a total of 30 cells per replicate. Cell selection was based on the acetylated α-tubulin signal to ensure unbiased identification of axonemal structures. Internalised c-rings were manually counted using the Cell Counter plugin in Fiji (ImageJ). All quantifications were performed using identical imaging and analysis settings across conditions. Statistical significance was calculated using one-way analysis of variance (ANOVA) and adjusted *p* values were derived using Šídá k’s multiple comparisons test and exact *p* values are included in the figure legend in Fig 1. *P* values were calculated using unpaired t test with Welch’s correction ****p<0.0001 in Fig. 4f.

Fluorescence intensity of tubulin post-translational modifications (PTMs) was quantified using Fiji (ImageJ). Regions of interest (ROIs) corresponding to external surface cilia or internalised axonemes/c-rings were selected based on acetylated α-tubulin signal, which was used as a reference marker for axonemal structures. Background ROIs were selected from adjacent regions lacking specific signal. Mean fluorescence intensities for PTMs and acetylated α-tubulin were measured using the ROI Manager. PTM enrichment was calculated as: *(PTM_ROI_-PTM_background_)/(Tubulin_ROI_−Tubulin_background_)*. All measurements were performed using identical acquisition and analysis settings across conditions. Data are presented as mean ± standard deviation (s.d.).

For quantification of newly regenerated cilia, five fields of view were selected at random for image acquisition of each biological replicate. The total number of cells in each field was counted by nuclear staining, and cells displaying new regenerated cilia were counted manually according to morphological criteria. The percentage of cells with regenerated cilia was calculated as: *Cells with new cilia (%) = (number of cells with new cilia/ total number of cells) * 100.* Data are presented as mean ± standard deviation (s.d.). *P* values were calculated using two-way analysis of variance (ANOVA) with Tukey’s multiple comparisons test. *P* values are included in the figure legend in Fig 5.

For quantification of cilia numbers, 10 cells were selected at random for each biological replicate. Cilia numbers were manually counted using the Cell Counter plugin in Fiji based on the z-projected images. Statistical significance was calculated using one-way analysis of variance (ANOVA) and adjusted *p* values were derived using Šáídk’s multiple comparisons test and exact *p* values are included in the figure legend in Fig 5.

### Proteomic analysis

#### Sample preparation

*Tetrahymena* cultures were grown in parallel in separate flasks for triplicates. When cultures reached an OD600 of 1.0, cells were transferred to 10 mM Tris-HCI and starved overnight. Cells were deciliated based on the above Calcium chloride based partial deciliation protocol. Following centrifugation, the cell pellets were collected, snap-frozen in liquid nitrogen, and stored until lysis.

#### TMT Labelling and High pH reversed-phase chromatography

Aliquots of 50 µg of each sample were digested with trypsin (1.25 µg trypsin; 37°C, overnight), labelled with Tandem Mass Tag (TMTpro) sixteen plex reagents according to the manufacturer’s protocol (Thermo Fisher Scientific, Loughborough, LE11 5RG, UK) and the labelled samples pooled. The pooled sample was desalted using a SepPak cartridge according to the manufacturer’s instructions (Waters, Milford, Massachusetts, USA). Eluate from the SepPak cartridge was evaporated to dryness and resuspended in buffer A (20 mM ammonium hydroxide, pH 10) prior to fractionation by high pH reversed-phase chromatography using an Ultimate 3000 liquid chromatography system (Thermo Fisher Scientific). In brief, the sample was loaded onto an XBridge BEH C18 Column (130Å, 3.5 µm, 2.1 mm x 150 mm, Waters, UK) in buffer A and peptides eluted with an increasing gradient of buffer B (20 mM Ammonium Hydroxide in acetonitrile, pH 10) from 0-95% over 60 minutes. The resulting fractions (concatenated into 10 in total) were evaporated to dryness and resuspended in 1% formic acid prior to analysis by nano-LC MSMS using an Orbitrap Fusion Lumos mass spectrometer (Thermo Scientific).

#### Nano-LC Mass Spectrometry

High pH RP fractions were further fractionated using an Ultimate 3000 nano-LC system in line with an Orbitrap Fusion Lumos mass spectrometer (Thermo Scientific). In brief, peptides in 1% (vol/vol) formic acid were injected onto an Acclaim PepMap C18 nano-trap column (Thermo Scientific). After washing with 0.5% (vol/vol) acetonitrile 0.1% (vol/vol) formic acid peptides were resolved on a 500 mm × 75 μm Acclaim PepMap C18 reverse phase analytical column (Thermo Scientific) over a 150 minute organic gradient, using 7 gradient segments (1-6% solvent B over 1 minute, 6-15% B over 58 minutes, 15-32%B over 58 minutes, 32-40%B over 5 minutes, 40-90%B over 1 minute, held at 90%B for 6 minutes and then reduced to 1%B over 1 minute.) with a flow rate of 300 nl min^−1^. Solvent A was 0.1% formic acid, and Solvent B was aqueous 80% acetonitrile in 0.1% formic acid. Peptides were ionized by nano-electrospray ionization at 2.0kV using a stainless-steel emitter with an internal diameter of 30 μm (Thermo Scientific) and a capillary temperature of 300°C.

All spectra were acquired using an Orbitrap Fusion Lumos mass spectrometer controlled by Xcalibur 3.0 software (Thermo Scientific) and operated in data-dependent acquisition mode using an SPS-MS3 workflow. FTMS1 spectra were collected at a resolution of 120 000, with an automatic gain control (AGC) target of 200 000 and a max injection time of 50ms. Precursors were filtered with an intensity threshold of 5000, according to charge state (to include charge states 2-7) and with monoisotopic peak determination set to Peptide. Previously interrogated precursors were excluded using a dynamic window (60s +/−10ppm). The MS2 precursors were isolated with a quadrupole isolation window of 0.7m/z. ITMS2 spectra were collected with an AGC target of 10,000, max injection time of 70ms and CID collision energy of 35%. For FTMS3 analysis, the Orbitrap was operated at 50,000 resolutions with an AGC target of 50 000 and a max injection time of 105ms. Precursors were fragmented by high energy collision dissociation (HCD) at a normalised collision energy of 60% to ensure maximal TMT reporter ion yield. Synchronous Precursor Selection (SPS) was enabled to include up to 10 MS2 fragment ions in the FTMS3 scan.

#### Data Analysis

The raw data files were processed and quantified using Proteome Discoverer software v2.4 (Thermo Scientific) and searched against the UniProt *Tetrahymena thermophila* database (downloaded August 2025: 26975 entries) using the SEQUEST HT algorithm. Peptide precursor mass tolerance was set at 10ppm, and MS/MS tolerance was set at 0.6Da. Search criteria included oxidation of methionine (+15.995Da), acetylation of the protein N-terminus (+42.011Da), methionine loss from the protein N-terminus (−131.04Da) and methionine loss plus acetylation of the protein N-terminus (−89.03Da) as variable modifications and carbamidomethylation of cysteine (+57.0214) and the addition of the TMTpro mass tag (+304.207) to peptide N-termini and lysine as fixed modifications. Searches were performed with full tryptic digestion and a maximum of 2 missed cleavages were allowed. The reverse database search option was enabled, and all data was filtered to satisfy false discovery rate (FDR) of 5%.

#### Proteomics data processing

Whole cell proteomic datasets were processed in MS Excel. Protein abundances for each of the three replicate runs were used to rank proteins by log_2_(foldchange) upregulation in levels at the 120-minute post-partial deciliation timepoint compared to the ciliated control samples. Two-tailed, unpaired t-tests assuming equal variances were performed to obtain an unadjusted *p* value for each protein from triplicate datasets for each dataset of samples to rank proteins consistently going up at 120-minutes post-partial deciliation samples compared to ciliated control samples (**Source Data Figure 3**). Proteins were visualised as individual dots in a scatter plot [log_2_(fold change) on the *x*-axis versus −log_10_*P* value on the *y*-axis] using GraphPad Prism v10. KEGG pathway enrichment analysis for each TMT dataset was performed using the ShinyGO 8.6 online tool (https://bioinformatics.sdstate.edu/go/). Most significantly upregulated top hits for each dataset were run against a background list of all *Tetrahymena* proteins detected in each TMT proteomic experiment to uncover enriched pathways using an FDR cutoff of 0.05.

#### Transmission electron microscopy

*Tetrahymena* cells were first fixed in 2.5% glutaraldehyde and 4% PFA in 0.1 M PIPES buffer, pH 7.2 for 1 hour at room temperature, then stored at 4°C. Cells were washed with 0.1 M PIPES buffer five times, followed by a 15-minute incubation in 50 mM glycine in buffer and one final wash in PIPES buffer only. Suspension cells were then enrobed in agarose to avoid the need for centrifugation. Cells underwent secondary fixation in 1% osmium tetroxide and 1.5% potassium ferrocyanide in buffer for 1 hour at 4°C, followed by washing with Milli-Q water. Tertiary fixation was performed in 0.5% uranyl acetate (aq.) overnight at 4°C. Cells were then dehydrated through a graded ethanol series (30%-100%), followed by resin infiltration (TAAB low viscosity) using a gradient of ethanol and resin mixtures. Cells were embedded in fresh resin and polymerized at 60°C for 24 hours. Ultrathin sections (∼90 nm) were cut using a DiATOME diamond knife on a Leica UC7 ultramicrotome and mounted onto formvar-coated grids. Sections were post-stained with lead citrate and imaged using a JEOL JEM-1400Flash 120kV transmission electron microscope equipped with a Gatan Rio camera. All reagents and equipment are provided by the Dunn School Electron Microscopy facility.

#### Volumetric electron microscopy

*Tetrahymena* cells were fixed, treated with 50 mM glycine and enrobed in agarose as per the procedure outlined above. Cells were further processed according to a modified rOTO protocol. Briefly, cells were post-fixed in 1.5% potassium ferricyanide and 2% osmium tetroxide in 0.1M PIPES buffer in the cold room for 1 hour, followed by washing with Milli-Q water. Cells were then treated with thiocarbonhydrazide (TCH) solution for 20 minutes on the rotator at room temperature, followed again by washing with distilled Milli-Q water. Cells were further fixed with 2% osmium tetroxide in Milli-Q water for 30 minutes on the rotator at 4℃. Tertiary fixation was performed in 1% uranyl acetate (aq.) at 4℃ overnight. Cells were then stained with lead aspartate for 30 minutes at room temperature on a rotator, followed by washing with Milli-Q water. Cells were then dehydrated through a graded ethanol series (30%-100%), followed by Durcupan resin infiltration using a series of ethanol and resin mixtures. Cells were embedded in fresh resin and polymerized at 60°C for 48-72 hours.

Polymerized sample blocks were trimmed by hand using a fine razor blade and mounted onto aluminium pins (Micro to Nano cat.10-006003) using conductive epoxy (Circuitworks cat. CW2400). Mounted samples were polished using a Leica UC7 ultramicrotome and sputter coated with 12 nm gold-palladium. Volume data were collected using a Zeiss Merlin VP compact with Gatan 3View2XP using an OnPoint back-scattered electron detector. The slice thickness was set to 50 nm. Images were acquired at 1.8keV (with focal charge compensation) with a frame size of 11,000 x 11,000 pixels (9 nm/pixel) and 1 µs dwell time. All data were processed using Dragonfly 3D world (v2025.1).

## Disclosure of generative AI use

OpenAI ChatGPT (GPT-6), accessed through the University of Oxford’s ChatGPT service, was used solely for language editing, formatting and refinement of the manuscript and cover letter. OpenAI Codex was used to develop and verify workflows for microscopy-video analyses. The manuscript was written and edited by the authors and reviewed by multiple human peers followed by further refinement and editing. All scientific content, interpretations, analyses, and conclusions were developed and verified by the authors, who take full responsibility for the submitted work.

## Acknowledgements

We thank Prof. Mark Winey and Dr. Rachel Anne-Howard Till for providing the RIB72B-mCherry strain and Prof. Doug Chalker and staff at the Tetrahymena Stock Center for providing the VPS13A-KO and VPS13A-GFP strains. We thank Dr. Alan Wainmann, Raman Dhaliwal and Dr. Jazmin Meza Torres for technical assistance with the SoRa spinning disk live cell microscopy, transmission electron microscopy and brightfield imaging respectively. We thank Profs. Shona Murphy, Sumana Sanyal, Drs. Georgia Isom, Thomas Williams, Aakash Mukhopadhyay, Alex Carver, Shiza Shaikh and Ms. Siham Abdul Jaleel for helpful comments on the manuscript. The authors would like to acknowledge Mr. Bradley Burnet for his preliminary observations of internalised ciliary c-rings in *Tetrahymena*.

## Author Contributions

M.R. optimized partial deciliation methodology, prepared all samples, acquired and formally analysed all data with help from C.M. for transmission and volumetric electron microscopy and segmentation with model building. K.H. and G.R.M performed TMT labelling and quantitative proteomic dataset acquisition and formal data analysis respectively. S.M. and K.N. provided the EGFP-TtATG8B strain. G.R.M conceived, supervised and acquired funding for the project. M.R. and G.R.M. prepared all figures, wrote the original draft and revised the manuscript with input from all authors.

## Data availability

All data are available in the main text or the supplementary materials. All mass spectrometry proteomics data have been deposited to the ProteomeXchange Consortium via the PRIDE partner repository under the dataset identifier PXD085083. The data are securely archived and available to editors and reviewers during the evaluation phase using confidential review credentials provided to the editor during submission. Upon acceptance of the manuscript, these data will be made public.

